# *Gpr63* is a novel modifier of microcephaly in *Ttc21b* mouse mutants

**DOI:** 10.1101/424341

**Authors:** J. Snedeker, WJ Gibbons, D.R. Prows, R.W. Stottmann

**Affiliations:** Division of Human Genetics, Cincinnati Children’s Hospital Medical Center University of Cincinnati College of Medicine, Cincinnati, OH; Department of Pediatrics, Cincinnati Children’s Hospital Medical Center University of Cincinnati College of Medicine, Cincinnati, OH; Neuroscience Graduate Program, Cincinnati Children’s Hospital Medical Center University of Cincinnati College of Medicine, Cincinnati, OH; Division of Developmental Biology, Cincinnati Children’s Hospital Medical Center University of Cincinnati College of Medicine, Cincinnati, OH

**Keywords:** Mouse, Quantitative trait locus, Microcephaly, Ttc21b, Primary cilia, Gpr63

## Abstract

The primary cilium is a critical signaling center for proper embryonic development. Previous studies have demonstrated that mice lacking *Ttc21b* have impaired retrograde trafficking within the cilium and multiple organogenesis phenotypes, including microcephaly. Interestingly, the severity of the microcephaly in *Ttc21b^aln/aln^* homozygous null mutants is considerably affected by the genetic background. *Ttc21b^aln/aln^* mutants on an FVB/NJ background develop a forebrain significantly smaller than mutants on a C57BL/6J background. We performed a Quantitative Trait Locus (QTL) analysis to identify potential genetic modifiers and identified two regions linked to differential forebrain size: *modifier of alien QTL1 (Moaq1)* on chromosome 4 at 27.8 Mb and *Moaq2* on chromosome 6 at 93.6 Mb. These QTLs were validated by constructing congenic strains. Further analysis of *Moaq1* identified a brain specific orphan G-protein coupled receptor (GPCR), *Gpr63*, as a candidate gene. We identified a SNP between the FVB and B6 strains in *Gpr63*, which creates a missense mutation predicted to be deleterious in the FVB protein. We first demonstrated that *Gpr63* can localize to the cilium and then used CRISPR-Cas9 genome editing to create FVB congenic mice with the B6 sequence of *Gpr63* and a deletion allele leading to a truncation of the GPR63 C-terminal tail. These alleles genetically interact with *Ttc21b^aln/aln^*, validating *Gpr63* as a forebrain modifier of *Ttc21b* and strongly supporting *Gpr63* as the variant causal gene (i.e., the quantitative trait gene, QTG) for *Moaq1*.

## INTRODUCTION

Primary cilia are microtubule-based organelles known to play essential roles in proper development and function of a number of organ systems including the central nervous system (CNS)(Guemez-Gamboa et al. 2014; Valente et al. 2014). Ciliopathies are a class of human diseases caused by ciliary mutations affecting the proper function of primary cilia (Goetz and Anderson 2010; Hildebrandt et al. 2011). The frequent presentation of cognitive impairment in ciliopathy patients in addition to severely compromised brain development in a number of ciliary mutant mouse models clearly displays the importance of primary cilia in CNS health and development (Han and Alvarez-Buylla 2010; Reiter and Leroux 2017).

Primary cilia have been implicated in transducing and regulating several critical developmental pathways including SHH, WNT, PDGF, TGFβ/BMP, RTK, and Notch (Goetz and Anderson 2010; Wheway et al. 2018). An array of cell surface receptors localize to the ciliary membrane to modulate these pathways and other signaling events and GPCRs represent a significant class of these ciliary receptors (Schou et al. 2015). Intraflagellar transport (IFT) proteins are responsible for the movement of cargo within the cilium with complex-B proteins regulating anterograde transport from the centriole to the distal tip of the cilium and complex-A proteins regulating retrograde transport (Prevo et al. 2017). IFT-A proteins have also been shown to specifically play a role in the trafficking of select GPCRs in and out of the cilium (Mukhopadhyay et al. 2010; Fu et al. 2016; Hirano et al. 2017).

*Tetratricopeptide repeat domain 21b (Ttc21b)* encodes a known IFT-A protein necessary for the proper rate of retrograde trafficking within the primary cilium (Tran et al. 2008). Proper ciliary trafficking is known to be critical for the processing of GLI transcription factors, which are in turn essential for proper regulation of the SHH pathway (Huangfu and Anderson 2005; Hui and Angers 2011). Mutant mice homozygous for the *alien* null allele of *Ttc21b* display impaired processing of GLI3, the primary repressor of SHH target genes, and consequently show an increase in SHH pathway activity in multiple tissues including the developing forebrain (Tran et al. 2008; Stottmann et al. 2009).

Genetic backgrounds can affect phenotypic severity and penetrance in mouse mutants, and these differences can be used to detect genetic interactions in complex traits (Doetschman 2009; Mackay 2014). This is true for the forebrain malformations seen in *Ttc21b^aln/aln^* mutants which presents an opportunity to identify modifying loci to help understand the underlying pathogenic molecular mechanism. Here we present such a QTL analysis and demonstrate that *Gpr63* interacts with *Ttc21b* to modify the size of the forebrain in *Ttc21b^aln/aln^* null mutants.

## RESULTS

### *Ttc21b^aln^* Microcephaly is Background Dependent

The microcephaly phenotype observed in *Ttc21b^aln/aln^* mutants was first identified as part of an ENU mutagenesis experiment in which the ENU mutagen was given to A/J mice. The subsequent breeding and mapping of the causative mutation involved an outcross to the FVB/NJ (FVB) strain (Herron et al. 2002; Tran et al. 2008). *Ttc21b^aln/aln^* homozygous mutants on this mixed A/J;FVB background exhibited multiple forebrain and craniofacial phenotypes. While the microcephaly phenotype was completely penetrant, craniofacial phenotypes, including cleft lip and palate, were only present in a minority of embryos. To increase the frequency of craniofacial phenotypes to facilitate future molecular analyses, the *Ttc21b^aln^* allele was serially backcrossed onto the C57BL/6J (B6) strain. The B6 genetic background has previously shown to be more susceptible to craniofacial phenotypes in a number of studies (Hide et al. 2002; Dixon and Dixon 2004; Mukhopadhyay et al. 2012; Percival et al. 2017). As the *Ttc21b^aln^* was serially backcrossed to the B6 background, we noticed the relative size of the forebrain tissues in B6.Cg-*Ttc21b^aln/aln^* mutants increased substantially, although they remained smaller than control brains and still lacked olfactory bulbs (Fig.1 E, F). To confirm this was indeed an effect of varying the genetic background, the *Ttc21b^aln^* allele was again serially backcrossed to the FVB background and the severity of microcephaly seen initially was recovered in FVB.Cg-*Ttc21b^aln/aln^* mutants (Fig. 1C, D). We measured the forebrain surface area of a number of *Ttc21b^aln/aln^* mutants on both FVB and B6 genetic backgrounds and saw a 60% decrease in forebrain size in FVB.Cg-*Ttc21b^aln/aln^* animals as compared to littermate controls. We observed a 30% decrease in B6.Cg-*Ttc21b^aln/aln^* animals. The interaction between genetic background and the *alien* mutation was found to be significant with a 2-way ANOVA (p=0.0114) and each individual wild-type; mutant pair was found to be statistically significant with a Student’s t-test (FVB, p=0.000054; B6, p=0.00033).

**Figure 1.**
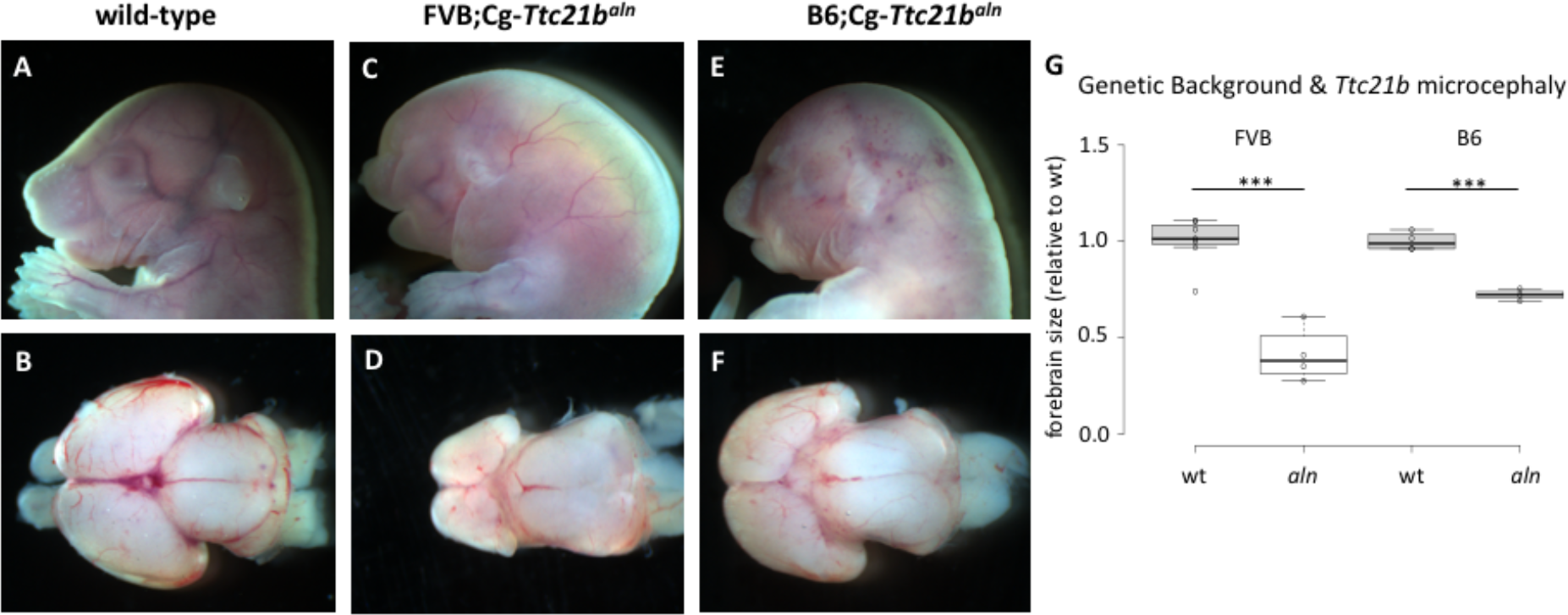
Genetic background affects the *Ttc21b^aln/aln^* microcephaly phenotype. (A-F) E18.5 embryos and whole mount brains viewed from the dorsal aspect from control (A, B), FVB;Cg-*Ttc21b^aln/aln^* mutants (C, D), and B6;Cg***-**Ttc21b^aln/aln^* mutants (E, F). (G) Relative brain sizes between mutants and wild-types for each genetic background. Center lines show the medians; box limits indicate the 25th and 75th percentiles as determined by R software; whiskers extend 1.5 times the interquartile range from the 25th and 75th percentiles, outliers are represented by dots. n = 7, 4, 4, 3 sample points. (*** p<0.005).

Independent of the genetic background, we also noted an incompletely penetrant exencephaly phenotype in *Ttc21b^aln/aln^* animals, but the frequency did not significantly differ between backgrounds. After a few generations of backcrossing onto FVB (FVB;B6-*Ttc21b^aln/aln^*) and B6 (B6;FVB-*Ttc21b^aln/aln^*), we performed a genome-wide high-density SNP scan and confirmed that each strain was at least 90% pure (Supplemental Table 1). We calculated the incidence of exencephaly and found that maintaining the *Ttc21b^aln^* allele on either the FVB or B6 genetic backgrounds yielded exencephaly in ~40% of *Ttc21b^aln/aln^* mutants (24/58, 41% in FVB;B6-*Ttc21b^aln^/aln* and 30/65, 46% in B6;FVB-*Ttc21b^aln^/aln*).

We hypothesized the genetic background effect on *Ttc21b^aln/aln^* mutant forebrain size (Fig. 1) would allow a Quantitative Trait Locus (QTL) analysis to shed insight into the underlying molecular mechanism(s). To maximize the benefit of a QTL experiment, we further purified the FVB and B6 advanced backcross lines with additional backcrosses combined with targeted microsatellite marker screening of chromosomal areas not homozygous for FVB or B6, as identified in the initial genome scan. This allowed us to create FVB.Cg-*Ttc21b^aln^* animals that were >99% FVB and B6.Cg-*Ttc21b^aln^* animals purified to ~97% B6 background, as assayed by a GigaMUGA genome SNP scan providing strain-specific sequence information at over 143,000 SNPs (Morgan et al. 2015) (Supplemental Table 1).

We first generated F2 *Ttc21b^aln/aln^* mutant progeny from a B6.FVB F1 (*Ttc21b^aln/+^)* intercross for QTL analysis. Fifty-four F2 litters were generated, which identified 148 *Ttc21b^aln/aln^* mutant embryos from 607 total embryos. We excluded 35 mutants (24%) with exencephaly, which precluded a measurement of forebrain area. Ninety-six brains from the remaining F2 mutants were microdissected and the dorsal surface area of the forebrain was measured. Whole genome SNP genotyping of each animal was carried out using a GigaMUGA panel and QTL mapping was performed using R/qtl (Broman and Sen 2009) to identify chromosomal regions linked to differential brain sizes in *Ttc21b^aln/aln^* mutants (Table S2).

A significant (genomewide p < 0.05) QTL (LOD 6.1) was mapped to chromosome 4 with its peak at 27.8 Mb (marker UNC6953268) and a 95% confidence interval spanning 19.5 Mb to 46 Mb (Fig. 2A, B). We have named this QTL *modifier of alien QTL 1 (Moaq1)*. A second QTL was identified with a LOD score suggesting significance (genomewide p < 0.63; LOD 4.7) and mapped to chromosome 6 at 93.6 Mb (UNCHS018175) with a 95% confidence interval from 0 to 117.5 Mb (Fig. 2A, C). We have named this QTL *modifier of alien QTL 2 (Moaq2)*. A third peak was found on chromosome 2 at the *Ttc21b^aln^* locus, which likely represents the A/J passenger background on which the initial mutagenesis was performed (Fig. 2A).

**Figure 2.**
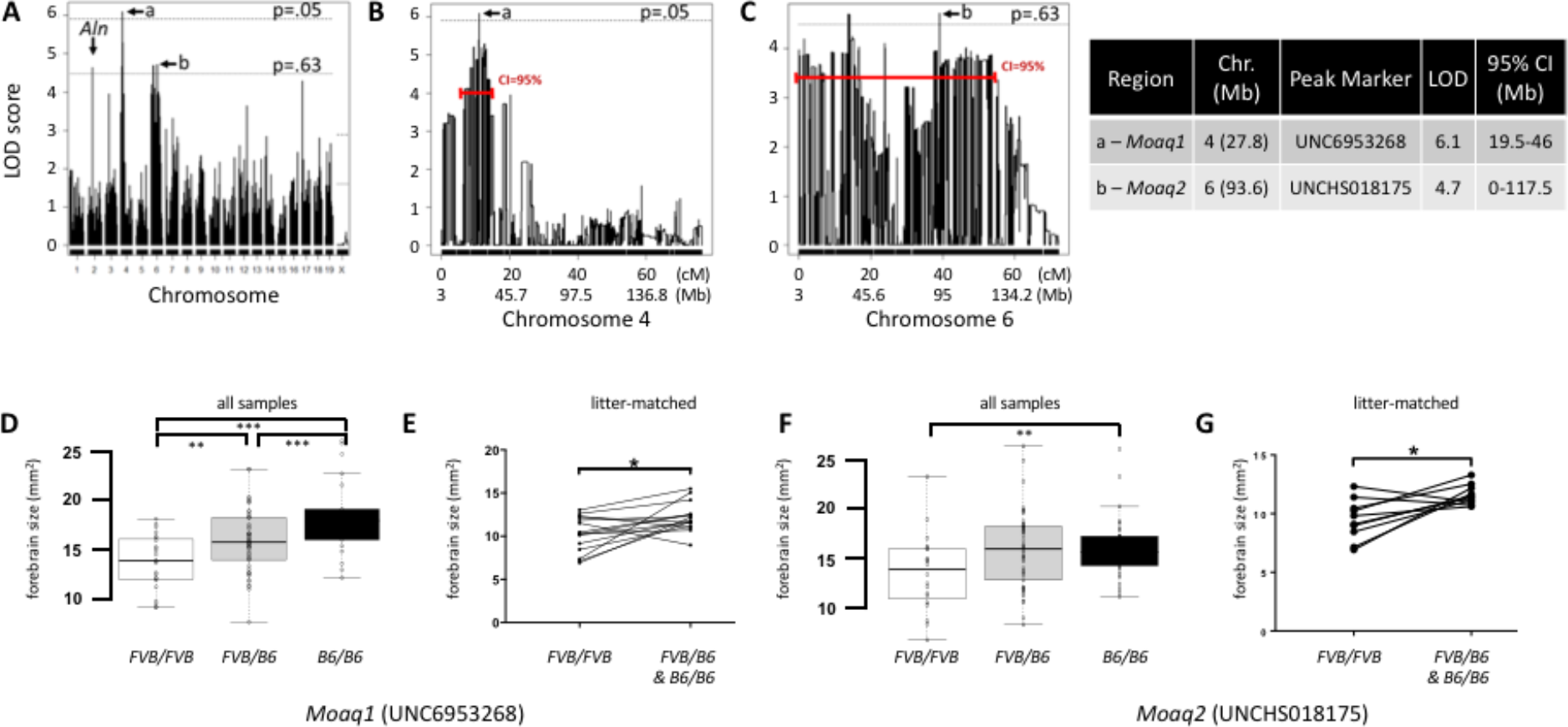
QTL analysis. (A) QTL LOD scores by chromosome with dotted lines displaying significance (genomewide p=0.05) and suggestive significance (genomewide p=0.63). (a) marks the significant QTL peak on chromosome 4, *Moaq1*, and (b) marks the suggestive *Moaq2* peak on chromosome 6. (B) Chromosome 4 QTL LOD scores and 95% confidence interval shown in red. (C) Chromosome 6 LOD scores and 95% confidence interval. (D) Analysis of brain size in animals homozygous for FVB genome identify, heterozygous for FVB/B6 and homozygous for B6 genome as genotyped by the peak marker for *Moaq1*. n = 24, 51, 21 sample points. (E) Analysis of brain size only among littermates in litters with both *Ttc21b^aln/aln^-Moaq1^FVB/FVB^* and *Ttc21b^aln/aln-^Moaq1^FVB/B6 <u>or</u> B6/B6^* embryos (n=15 litters). (F) Analysis of brain size in animals homozygous for FVB genome identify, heterozygous for FVB/B6 and homozygous for B6 genome as genotyped by the peak marker for *Moaq2*. n = 24, 51, 21 sample points. (G) Analysis of brain size only among littermates in litters with both *Ttc21b^aln/aln^-Moaq2^FVB/FVB^* and *Ttc21b^aln/aln^-Moaq2^FVB/B6 <u>or</u> B6/B6^* embryos (n=6 litters). (*** p<0.005; ** p<0.05, * p≤0.01)

We confirmed a correlation between the genetic identity of *Moaq1* and *Moaq2* and an effect on forebrain size by comparing the genotype of the peak SNP marker for each QTL with forebrain size (Fig. 2D, F). We first independently assessed F2 mice for genotype at the peak marker of the *Moaq1* locus and found an incremental decrease in brain size among *Ttc21b^aln^/aln-Moaq1^FVB/FVB^* animals as compared to *Ttc21b^aln^/aln-Moaq1^FVB/B6^* or *Ttc21b^aln^/aln-Moaq1^B6/B6^* animals (Fig. 2D). An overall ANOVA indicated a statistically significant difference (p<0.001) and a subsequent Tukey analysis showed each group was significantly different (p<0.001). To control for a potential litter-based difference, we also compared *Ttc21b^aln/aln^-Moaq1^FVB/FVB^* mutants with *Ttc21b^aln/aln^-Moaq1^FVB/B6^* or *Ttc21b^aln/aln^-Moaq1^B6/B6^* littermates (Fig. 2E). Although this analysis had less statistical power, we again saw a significant reduction in *Ttc21b^aln/aln^-Moaq1^FVB/FVB^* animals (a 1.94mm^2^ decrease; p=0.012).

A similar analysis testing just the *Moaq2* locus showed a significant variation among *Ttc21b^aln/aln^-Moaq2^FVB/FVB^*, *Ttc21b^aln/aln^-Moaq2^FVB/B6^* and *Ttc21b^aln/aln^-Moaq2^B6/B6^* animals (Fig. 2F, p=0.0243). However, a subsequent analysis only found a significant individual difference between *Ttc21b^aln^/aln-Moaq2^FVB/FVB^* and *Ttc21b^aln/aln^-Moaq2^B6/B6^* (p=0.0255). We again complemented this with a littermate only analysis and found *Ttc21b^aln/aln^-Moaq2^FVB/FVB^* animals had approximately 20% smaller forebrains than littermates (Fig. 2G; p=0.01).

### Congenic Lines

We produced independent congenics to replicate the *Moaq1* and *Moaq2* loci. We noted in our original F2 population of 96 mice for the QTL analysis no *Ttc21b^aln/aln^* mutants were homozygous for B6 SNPs over a 10-Mb region on chromosome 14. We hypothesized this represented a region of the B6 genome that, in combination with homozygosity for the *Ttc21b^aln^* allele, leads to either early embryonic lethality and/or exencephaly. Thus, we attempted to create FVB congenics with *Moaq1* and *Moaq2* genomic loci from the B6 strain. B6;FVB-*Ttc21b^aln^/wt* F1 mice, heterozygous across their genome, were repetitively backcrossed with FVB mice to produce a congenic strain that was FVB background except for the heterozygous *Moaq1* or *Moaq2* B6 QTLs (*i.e*., FVB.Cg-*Moaq1^FVB/B6^* and FVB.Cg-*Moaq2^FVB/B6^*). Mice from each congenic line were then intercrossed to determine if the B6-derived *Moaq1* and/or *Moaq2* QTLs could influence forebrain size on their own. As we analyzed the genomes of FVB.Cg-*Ttc21b^aln^/aln* mutants at the end-stage of congenic construction, we observed *Ttc21b^aln^/aln* mutants on a >99% FVB background all either died prior to E17.5 or had exencephaly precluding measurement of brain size (more than 50 FVB.Cg-*Ttc21b^aln/aln^* mutants were produced with unusable brains). This suggests that the B6 genomic regions lost during the later backcrosses (i.e., as we progressed from 90 to 99% FVB) have a major influence on the incidence of exencephaly. Given that the increasing incidence of exencephaly precluded an analysis of forebrain size in the congenic lines (N5 and greater), we analyzed animals from the earlier backcross generations. Littermate brains were analyzed using paired t-tests to account for potentially confounding variables, such as length of gestation and/or background differences based on the generation of backcross. The *Moaq1* QTL was found to significantly affect brain size in FVB;B6-*Ttc21b^aln/aln^* mutants in the N2 to N5 generations. *Ttc21b^aln/aln^-Moaq1^FVB/B6^* and *Ttc21b^aln^/aln-Moaq1^B6/B6^* animals had larger brains as compared to *Ttc21b^aln/aln^-Moaq1^FVB/FVB^* (Fig. 3A), with an average decrease of 2.24mm^2^ (Fig. 3B; p=0.0177). The *Moaq2* QTL was also validated using this strategy and *Ttc21b^aln/aln^-Moaq1^FVB/B6^* and *Ttc21b^aln/aln^-Moaq1^B6/B6^* animals again had larger brains than *Ttc21b^aln/aln^-Moaq1^FVB/FVB^* (Fig. 3C), with an average decrease of 3.40mm^2^ (Fig. 3D; p=0.020). Thus, we conclude the QTL analysis of just 96 B6;FVB-*Ttc21b^aln/aln^* F2 forebrains produced two validated, novel QTLs that modify the effect of the *Ttc21b^aln/aln^* mutation on forebrain size.

**Figure 3.**
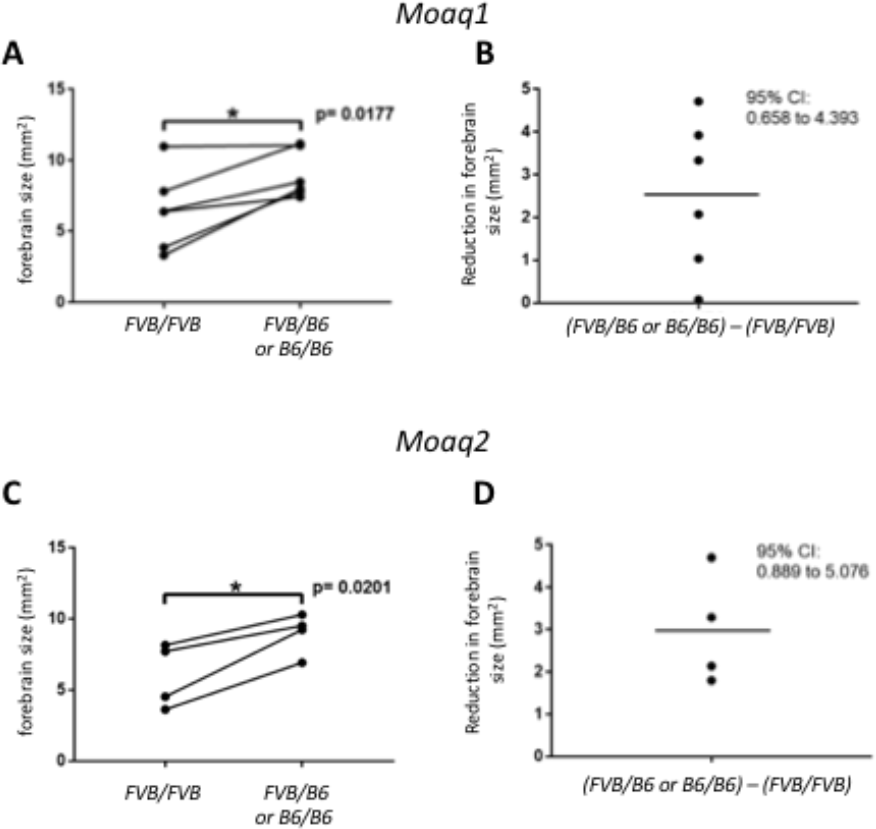
Congenic line validation of *Moaq1* and *Moaq2*. (A) Forebrain size from *Ttc21b^aln/aln-^Moaq1^FVB/FVB^* (FVB/FVB) and *Ttc21b^aln/aln^-Moaq1^FVB/B6 <u>or</u> B6/B6^* (FVB/B6 or B6/B6). (B) The amount of forebrain reduction in (FVB/B6 or B6/B6) – (FVB/FVB) brain sizes shows an average decrease of 2.24mm^2^. (C, D) Parallel analyses for *Moaq2* animals are shown. (* p≤0.02)

### Candidate Genes for *Moaq1* QTL

In order to identify potential candidate causal genes within the ~26.5 Mb *Moaq1* QTL, the 253 genes within the 95% confidence interval were analyzed (Table S3). We first identified 35 genes with missense variants when comparing B6 and FVB genomic sequences. The list of genes with insertions, deletions, and missense changes was further refined to 24 (Table S3) by selecting only those genes in which the A/J and FVB mutation were the same given the phenotypic similarities in *Ttc21b^aln/aln^* mutants on those genetic backgrounds. The remaining missense mutations were analyzed with the SIFT prediction algorithm (Kumar et al. 2009) and five genes were found with predicted deleterious missense mutations (<0.05 SIFT value). In addition to the 5 missense changes, 3 genes had indels which could not be analyzed by SIFT. However, the PROVEAN algorithm (Choi et al. 2012) made predictions of “neutral effect” for two of these and they were not included in the next analysis. We then performed a literature review of the 6 remaining genes (Table 1) for a plausible link to developmental neurobiology and narrowed the list to 3 candidates: *gamma-glutamyl hydrolase (Ggh), G-protein coupled receptor 63 (Gpr63)*, and *origin recognition complex subunit 3 (Orc3). Ggh* is a lysosomal enzyme with a role in folate metabolism which is provocative given the exencephaly noted in *Ttc21b^aln/aln^* mutants (Greene et al. 2009). Furthermore, *Ggh* is the only gene in the interval with 2 potentially deleterious missense mutations. *Orc3*, is a nuclear localized component of the Origin Recognition Complex (Bell 2002; Schneider and Ryan 2006). While conditional ablation of *Orc3* in the brain revealed a role in proper neural progenitor and radial glial development, germline loss of *Orc3* led to early embryonic lethality (Ma et al. 2013). A central role in DNA replication suggests a role for *Orc3* beyond neural development.

**Table 1.**
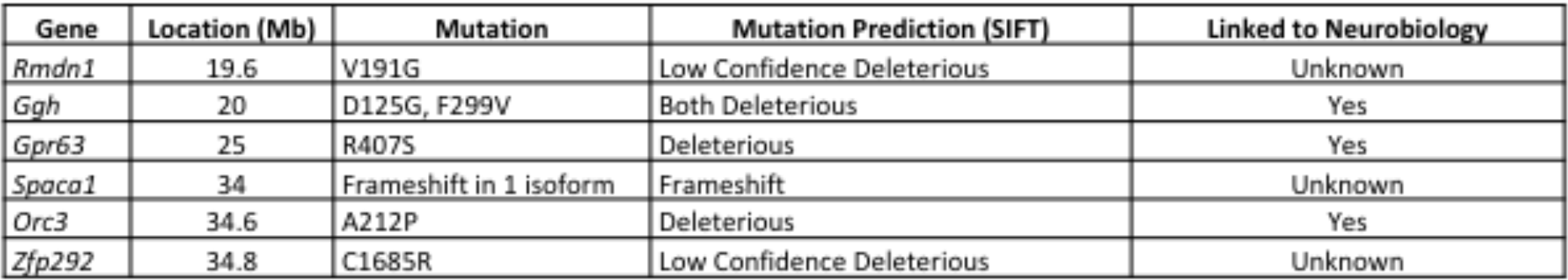
*Moaq1* candidate genes.

We next considered any evidence for candidate genes within the *Moaq1* QTL to have a role in primary cilia biology. Unlike *Ggh* or *Orc3, Gpr63*, is a membrane protein, making it the more attractive candidate to be a modifier of a mutation in a plasma membrane-based organelle, the primary cilium. *Gpr63* has been reported to have brain-specific expression and enriched in the forebrain (Kawasawa et al. 2000; Lee et al. 2001). The predicted deleterious polymorphic SNP within *Gpr63* (rs13477613; NM_030733 c.G1635T; NP_109658 p.R407S; SIFT score of 0.01) is a missense variant altering the coding of amino acid 407 from Arginine, an amino acid with an electrically charged side chain, in the B6 reference genome to Serine, an amino acid with a polar uncharged side chain, in the A/J and FVB genomes (Table 1). This Arginine residue is conserved from frogs to chimpanzee (Fig. 4.A), although humans have a Lysine at the orthologous residue in GPR63, an amino acid with an electrically charged side chain like Arginine. The loss of Arginine from the mouse GPR63 cytoplasmic tail alters a predicted RxR endoplasmic reticulum (ER) retention signal (Fig. 4A) (Michelsen et al. 2005). We directly tested the hypothesis that *Gpr63* might, at least partially, localize to the primary cilium by overexpressing a myc-tagged GPR63 construct in mouse Inner Medullary Collecting Duct (IMCD) cells. We chose IMCD cells as they are an accepted cellular model for ciliary biology and relatively easy to transfect. We do indeed see that *Gpr63-myc* co-localizes with acetylated tubulin marking the axoneme of the primary cilia (Fig. 4B-I). We note that not all the *Gpr63-myc* signal is in the primary cilium in a pattern consistent with non-ciliary plasma membrane. Taken together, we therefore hypothesized that the *Gpr63* polymorphism may be the causal variant within the *Moaq1* QTL.

**Figure 4.**
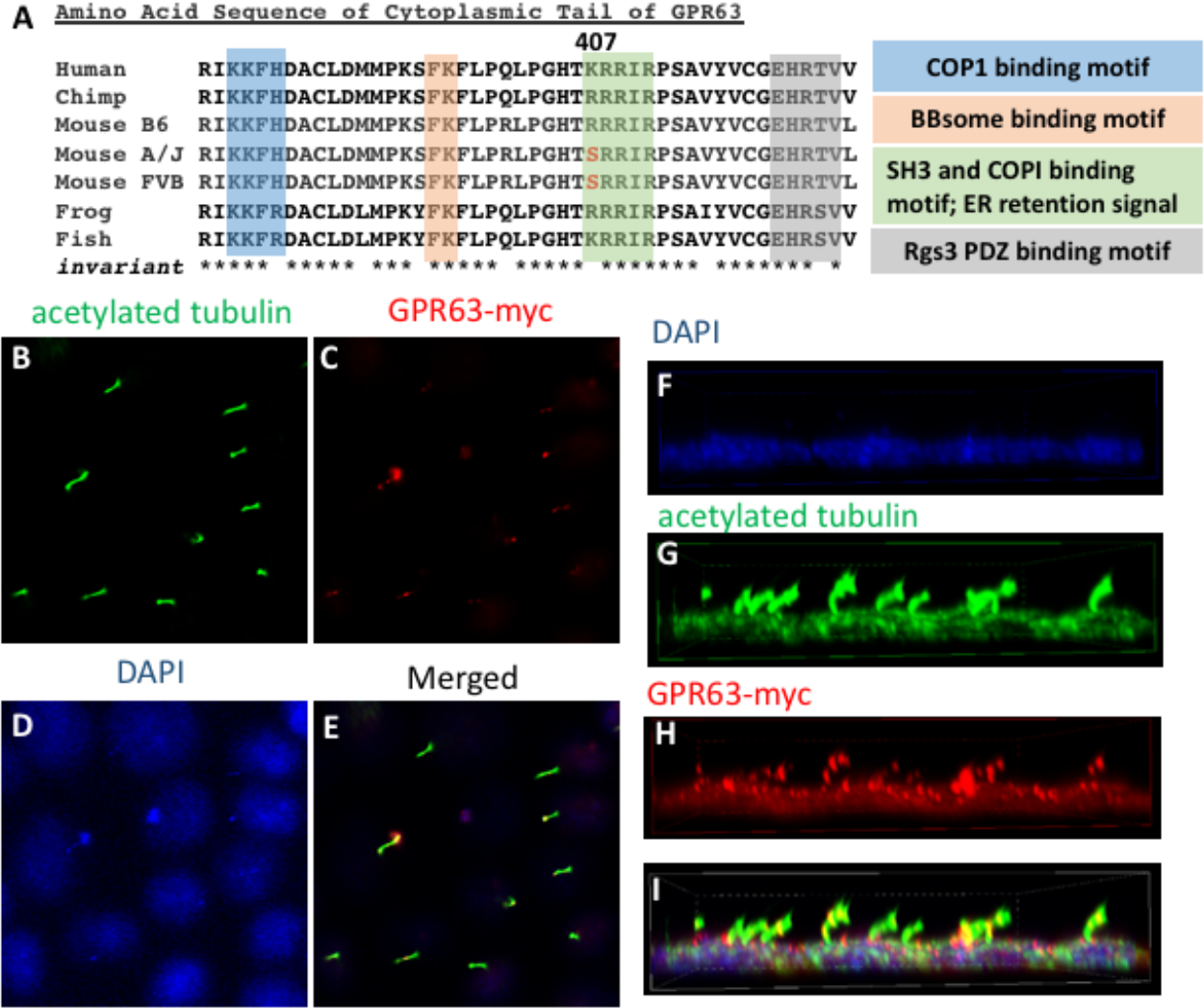
*Gpr63* is a conserved ciliary gene. (A) Amino acid sequence of GPR63 from multiple species and three inbred strains of mice. Invariant amino acids are indicated with *. Shading indicates amino acids composing four putative functional domains in the cytoplasmic tail of GPR63. (B-I) Overexpression of GPR63-myc *in vitro* leads to protein localization in the primary cilium. IMCD3 cells were transfected with *Gpr63-myc* plasmid and grown to confluency. Immunocytochemistry for acetylated tubulin to mark the axoneme (B), myc (C), DAPI for nuclei (D) shows some colocalization of *Gpr63-myc* and tubulin indicating ciliary localization (E = merged image). (F-I) A three-dimensional rendering of parallel experiment shows *Gpr63-myc* in both the cilium and plasma membrane.

### CRISPR-Cas9 Genome Editing of *Gpr63*

A standard approach to reduce the number of candidate genes within a QTL is to narrow the genetic interval carrying the trait of interest using meiotic recombination. For this, QTL-interval specific recombinants are identified and overlapping subcongenics produced and tested to determine the minimal region of effect. This process can be repeated, as needed. We initially pursued this strategy, but the results above (Fig. 3) show that the combination of exencephaly and early lethality of congenic FVB.Cg-*Ttc21b^aln/aln^* mice will preclude successful implementation unless we could also identify the region contributing to the exencephaly.

An alternative approach is to directly test hypotheses about candidate sequence variant(s) with genome editing to create novel transgenic models. We chose to pursue this and targeted FVB mice to change the rs13477613 polymorphism from FVB sequence (c.1635-T; coding for *Gpr63^Ser(FVB)^*) to the reference, non-pathogenic, B6 sequence (c.1635-G; coding for *Gpr63^Arg(B6)^*). This allele *Gpr63^em1Rstot^*, is hereafter referred to as FVB.Cg*-Gpr63^Arg(B6)^*. These animals were crossed with FVB.Cg-*Ttc21b^aln/wt^* (which will have the *FVB* allele of Gpr63: *Gpr63^Ser(FVB)^)* mice to produce FVB.Cg-*Ttc21b^aln/wt^*-*Gpr63^Arg(B6)^/Ser(FVB)* double heterozygotes. This was designed to directly test the hypothesis that the predicted pathogenic polymorphism of *Gpr63* contributes to the more severe microcephaly seen in FVB.Cg-*Ttc21b^aln/aln^* homozygous mutants. However, we were unable to directly test this hypothesis. As this experiment was done concurrently with the backcrosses to produce FVB congenic lines described above, we also found that FVB.Cg-*Ttc21b^aln/^aln*-*FVB.Gpr63^Arg(B6)/Arg(B6)^* mice have exencephaly and early lethality at a frequency precluding measurement of forebrain size in these animals as compared to control FVB.Cg-*Ttc21b^aln/aln^*-*FVB.Gpr63 Ser(FVB)*/*Ser(FVB)*.

An added benefit of CRISPR/Cas9 genome editing is the potential to recover an allelic series from the initial series of transgenic founders. We recovered and maintained a second allele *Gpr63^em2Rstot^* (*Gpr63^Del^*), with an 8-bp deletion resulting in a frameshift mutation leading to the translation of 5 missense amino acids and an early stop, preventing translation of the last 21 amino acids of the GPR63 C-terminus (Fig. 5A). This *Gpr63^Del^* mutation may disrupt both an ER retention signal and an Rg3 PDZ binding motif, creating a potentially more deleterious mutation than *Gpr63^Ser(FVB)^* (Michelsen et al. 2005; Saksela and Permi 2012; Kundu et al. 2013; Kundu and Backofen 2014; Kundu et al. 2014).

**Figure 5.**
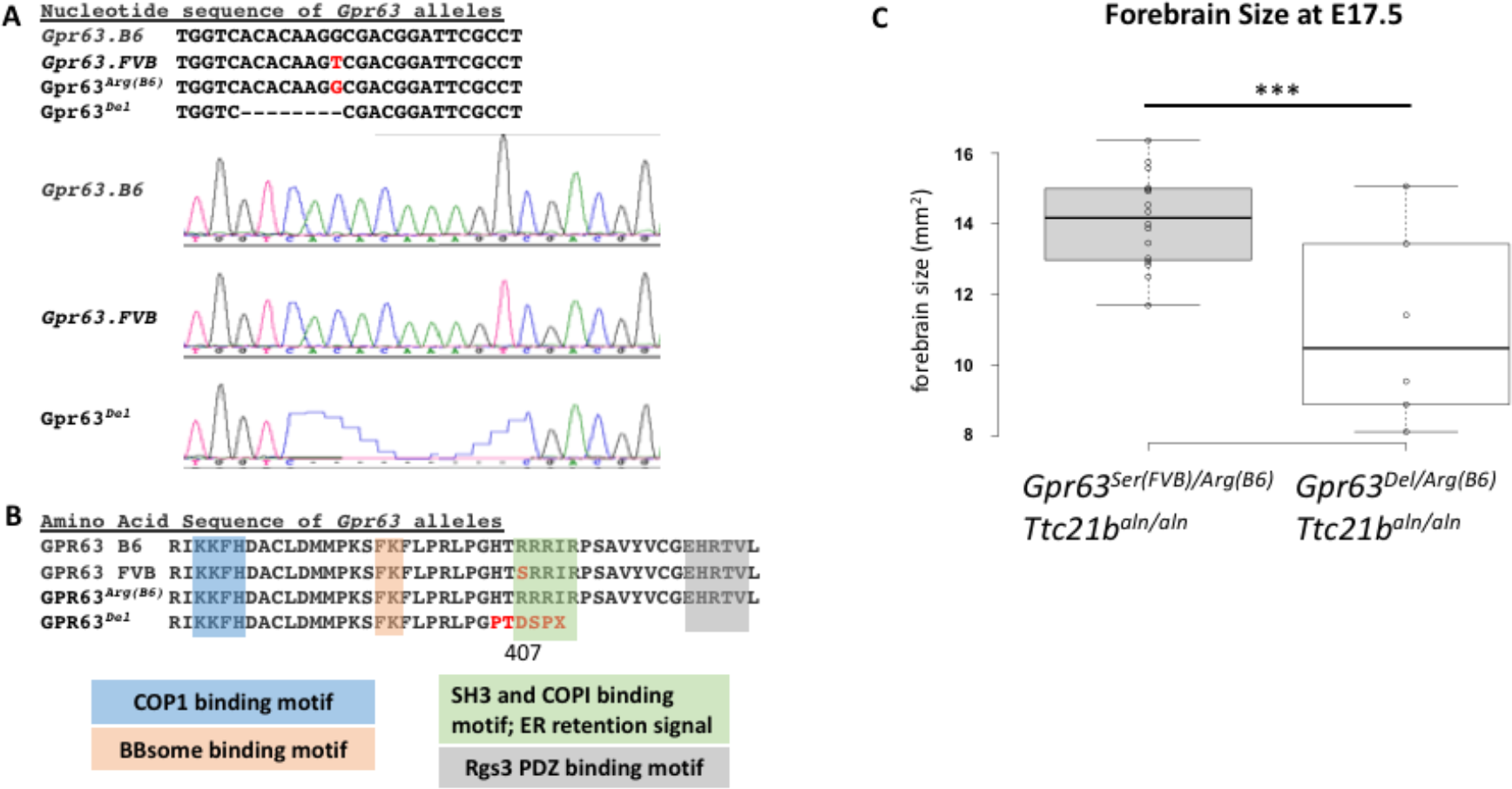
CRISPR-CAS9 alleles of *Gpr63* interact with *Ttc21b*. (A) Nucleotide sequence and Sanger sequencing of *Gpr63* in the B6/reference strain and FVB strain with the c.G1635T SNP. *Gpr63^Arg(B6)^* is the FVB mouse line with the B6 Thymidine at *Gpr63* c.1635. *Gpr63^Del^* is an 8-bp deletion in the same region of *Gpr63*. (B) Amino acid sequence of the above lines showing the R407S polymorphism between B6 (Arg) and FVB (Ser) reverted to B6 (Arg) in the CRISPR transgenic, and the missense mutation and premature truncation in the *Gpr63^Del^* allele. (C) Forebrain size in FVB;B6-*Ttc21b^aln/aln^-Gpr63^Arg(B6)/^Ser(FVB)* E17.5 embryos is significantly larger than FVB;B6-*Ttc21b^aln/aln^*-*Gpr63^Del/Arg(B6)^* animals. (*** p<0.005).

### *Gpr63* modifies forebrain phenotype in *Ttc21b^aln/aln^* mutants

We hypothesized the *Gpr63^Del^* allele may be more deleterious than the *Gpr63^Ser(FVB)^* polymorphism alone. We first determined if this allele affects survival and created animals homozygous for the *Gpr63^Del^* allele. We recovered *Gpr63^Del^/Del* mutants in approximately Mendelian ratios (5/16) and they survived to adulthood (n=5) with no discernible phenotypes seen to date. Consistent with the predicted deleterious SNP seen in both FVB and A/J animals, this suggests that *Gpr63* does not have an essential requirement in development.

In order to test for a genetic interaction with *Ttc21b*, we created FVB.Cg-*Ttc21b^aln/wt^-Gpr63^Del/Ser(FVB)^* mice and crossed these with B6.Cg-*Ttc21b^aln/wt^* (with no modification of the Gpr63 locus, e.g., B6.Cg-*Ttc21b^aln/wt^*-*Gpr63^Arg(B6)/Arg(B6)^* to compare F1 FVB;B6-*Ttc21b^aln/aln^-Gpr63 Ser(FVB)/Arg(B6)* with F1 FVB;B6-*Ttc21b^aln/aln^*-*Gpr63^Del/Arg(B6)^*. We hypothesized that reduced GPR63 function in the F1 FVB;B6-*Ttc21b^aln/aln^*-*Gpr63^)^Del/Arg(B6)* embryos as compared to F1 FVB;B6-*Ttc21b^aln/aln^-Gpr63 Ser(FVB/Arg(B6)* will lead to a smaller forebrain in the F1 FVB;B6-*Ttc21b^aln/aln-^Gpr63^Del/Arg(B6)^* animals. Indeed, F1 FVB;B6-*Ttc21b^aln/aln^-Gpr63^Del/Arg(B6)^* mice had a significant (~25%) decrease in brain size (Fig. 5B; −2.36mm^2^; n=15 and n=6; p=0.0019). This experiment demonstrates that *Gpr63* does indeed modify the *Ttc21b^aln/aln^* forebrain phenotype and supports the conclusion that *Gpr63* genetically interacts with *Ttc21b*.

To further address a possible interaction between *Ttc21b* and *Gpr63*, we aimed to create double homozygous null animals. We intercrossed F1 FVB;B6-*Ttc21b^aln/wt^-Gpr63^Del/Arg(B6)^* to compare F2 FVB;B6-*Ttc21b^aln/aln^-Gpr63^Arg(B6)/Arg(B6)^*, F2 FVB;B6-*Ttc21b^aln/aln^-Gpr63^Del/Arg(B6)^*, and F2 FVB;B6-*Ttc21b^aln/aln^-Gpr63^Del/Del^*. The F2 FVB;B6-*Ttc21b^aln/aln^-Gpr63^Del/Del^* animals are predicted to be more severely affected by loss or reduction of both *Ttc21b* and *Gpr63* function. We recovered 10 *Ttc21b^aln/aln^* embryos at E17.5 from this cross and found only slight reduction in forebrain size in F2 FVB;B6-*Ttc21b^aln/aln^-Gpr63^Del/Arg(B6)^* as compared to F2 FVB;B6-*Ttc21b^aln/aln-^Gpr63^Arg(B6)/Arg(B6)^* (Fig. 6). The small sample size does not allow us to make a rigorous conclusion with the data to date about this interaction (p=0.27). However, the most intriguing finding from this cross is that all three F2 FVB;B6-*Ttc21b^aln/aln^-Gpr63^Del/Del^* embryos recovered at E17.5 were necrotic and likely died around E12.5. Thus, we conclude that an allele of *Gpr63* with a truncation of the cytoplasmic tail interacts with *Ttc21b*, such that double mutants are embryonic lethal at stages prior to any lethality seen in either single mutant. The mechanism(s) leading to this lethality will be the subject of future study. We conclude from this study that *Gpr63* is likely the causal gene within the *Moaq1* QTL and reduced function of GPR63 in the FVB and A/J genetic backgrounds leads to the more severe forebrain phenotype of the *Ttc21b^aln/aln^* homozygous mutants on these backgrounds, as compared to those recovered on a B6 inbred background.

**Figure 6.**
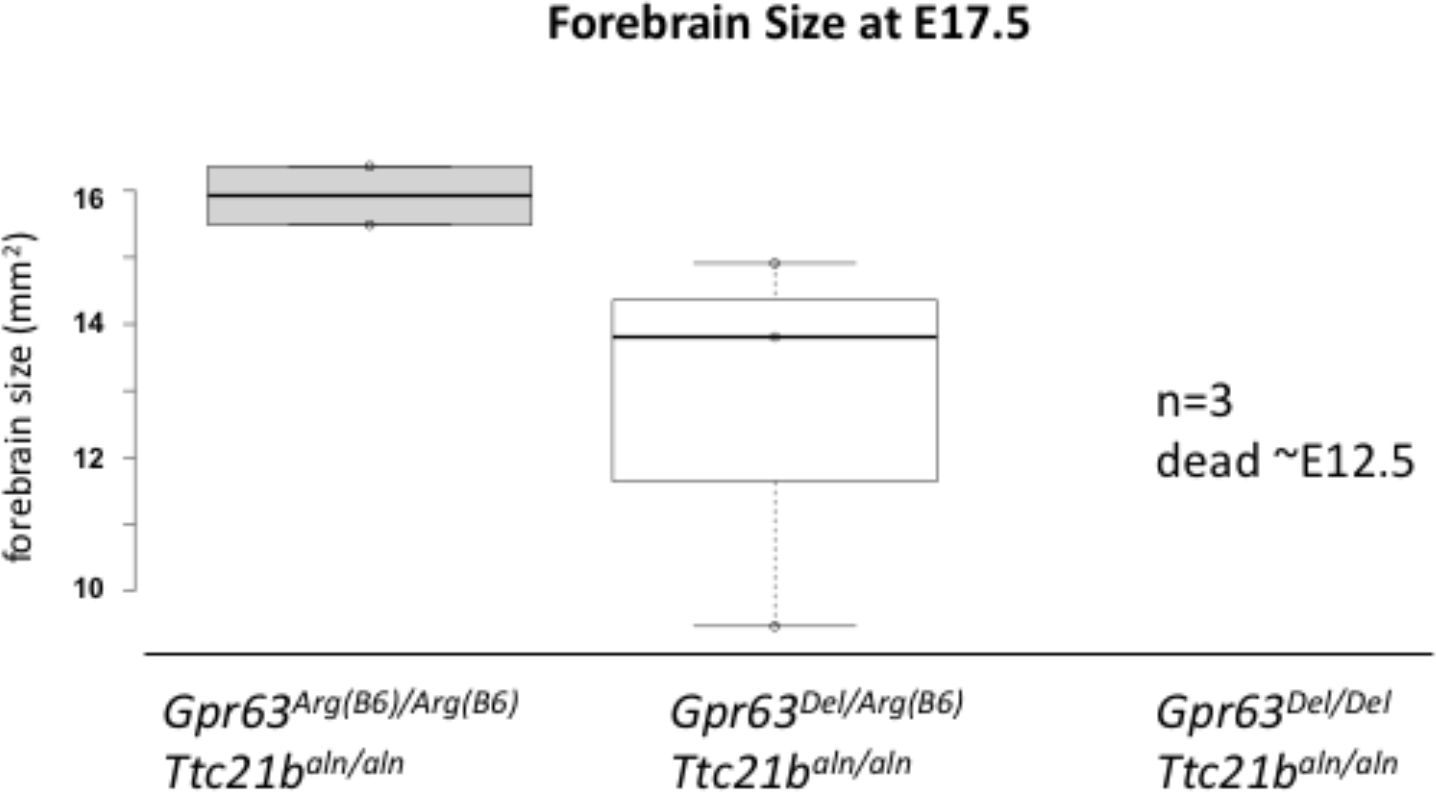
*Gpr63-Ttc21b* double mutants do not survive to term. Forebrain size in FVB;B6-*Ttc21b^aln/aln^-Gpr63^Arg(B6)/Arg(B6)^* animals is greater than FVB;B6-*Ttc21b^aln/aln^-Gpr63^Del/Arg(B6)^*. All three FVB;B6-*Ttc21b^aln/aln^-Gpr63^Del/Del^* animals recovered to date (n=3) had died well before E17.5.

## DISCUSSION

In this study we sought to understand the genetic basis for the influence of mouse inbred strain background on the severity of the *Ttc21b^aln/aln^* homozygous mutant microcephaly phenotype. Our QTL analysis identified two loci (*Moaq1, Moaq2)* which interact with *Ttc21b* to control brain size in *Ttc21b^aln/aln^* mutants and were validated by constructing QTL-containing congenic lines for each. The 95% confidence interval of *Moaq1* was small enough to allow a candidate gene approach and *Gpr63* was selected for further study. Experiments directly testing *Gpr63* function with novel mouse alleles generated with CRISPR/CAS-9 genome editing support our hypothesis that reduced *Gpr63* function further compromises embryos lacking functional *Ttc21b*. We also show that the *Ttc21b^aln^* allele is more susceptible to neural tube defects on the FVB genetic background, suggesting future identification of other genetic interactions may elucidate fundamental mechanisms of structural birth defects.

### Meiotic recombination or genome editing

Traditionally, the identification of a QTL is followed by a laborious mapping process and the creation of congenic strains to validate the QTL and begin to hunt for the truly causal sequence difference (Flint et al. 2005; Drinkwater and Gould 2012). The combination of deeply sequenced mouse inbred strains combined with the capabilities conferred by new genome editing tools such as CRISPR-CAS9 to rapidly generate sequence specific alterations in novel mouse lines can potentially greatly facilitate the identification of causal genetic variants within a QTL. In this case, we identified a potentially deleterious SNP in *Gpr63* on the FVB and A/J genetic backgrounds which may cause reduced function as compared to the canonical sequence seen in B6 mice. We continued backcrossing to FVB towards congenic line completion, but were confounded by the increased incidence of exencephaly and/or early embryonic lethality seen in *Ttc21b^aln/aln^* mutants on the >90% FVB genetic background. Presumably this increased frequency in exencephaly was due to the remaining 10% of the heterozygous genome that was further purified to FVB during this backcrossing, an important finding that could potentially facilitate mapping those modifiers independently. In parallel, we attempted to bypass the meiotic recombination and QTL refinement entirely by recreating the specific *Gpr63* SNP and determining if it affected brain size in the *Ttc21b* mutant background. No embryos without exencephaly and homozygous for B6 at a region on chromosome 14 were found in the initial QTL analysis. This suggested that the *Ttc21b^aln^* allele on a pure B6 background may be early embryonic lethal; therefore, we chose to do this by creating the B6 SNP on an FVB background (FVB *Gpr63^Arg(B6)/Ser(FVB)^*). Unfortunately, this has since been shown not to be the case. In contrast, the consequences of creating a pure inbred FVB strain with the *Ttc21b* allele prevented us from testing this *Gpr63* hypothesis directly, as pure FVB animals do have exencephaly and/or early lethality. However, the truncated *Gpr63* allele we recovered from our CRISPR-CAS9 studies did allow us to test this hypothesis and see an effect (Fig. 5). The best evidence for a genetic interaction between *Gpr63* and *Ttc21b* is the fact that all F2 FVB;B6-*Ttc21b^aln/aln^-Gpr63^Del/Del^* embryos recovered to date did <u>not</u> survive to E17.5: a phenotype not seen in either single mutant phenotype alone.

### Gpr63 and Ttc21b interactions

Having established a genetic interaction between *Gpr63* and *Ttc21b*, an important next step will be to determine if and how GPR63 and TTC21B physically interact. A direct biochemical interaction would significantly support the model. Our data suggest deleterious mutations in *Gpr63* are tolerated when normal ciliary function is occurring and that it contributes to more significant phenotypes when proper retrograde trafficking is impaired, such as in the context of *Ttc21b* mutations. One model to explain this set of results is if TTC21B were required for trafficking GPR63 out of the cilium, as it is known to do for other GPCRs (Hirano et al. 2017). If GPR63 function is impaired, perhaps it can be removed by TTC21B under normal conditions before significant negative effects accumulate. However, in the context of an animal lacking functional *Ttc21b*, it may remain in the cilium further perturbing developmentally important signal transduction. A careful study of GPR63 ciliary localization in both the presence and absence of TTC21B, and during dynamic developmental signaling, will be very informative.

The C-terminus of GPCRs is understood to play critical roles in protein-protein interactions and regulation of the receptor (Bockaert et al. 2003). As shown in Fig. 4, the c.G1635T SNP changes the amino acid sequence of the coat protein complex I (COPI) binding sequence. COPI is a coatomer responsible for the retrograde transport of vesicles from the *trans*-Golgi network to the *cis*-Golgi and from the *cis*-Golgi to the ER (Duden 2003; McMahon and Mills 2004). COPI has been implicated in the retrieval of proteins with exposed RxR motifs (Zerangue et al. 2001; Yuan et al. 2003). These motifs function to retain proteins in the ER until the RxR containing protein is completely processed and ready to be transported out through the Golgi (Michelsen et al. 2005). Often these proteins escape the RxR retrieval mechanism by masking the motif through interaction with another protein, a critical step in preparing the protein for its function which may include assembling a heteromultimeric complex (Michelsen et al. 2005). There are candidate motifs for protein domain interactions on the GPR63 C-tail which could serve in this masking role including: LPRLPGH, a predicted SH3 binding motif of DLG4, and EHRTV, a predicted PDZ binding domain of RGS3 (Saksela and Permi 2012; Kundu et al. 2013; Kundu and Backofen 2014; Kundu et al. 2014).

A defective RxR ER retrieval domain in GPR63 suggests 3 potential ways the protein could be impaired in FVB animals: excessive GPR63 localization to the cilium, premature exit of GPR63 from ER before it is properly processed, or a combination of the two resulting in excessive localization of improperly processed GPR63. In each case, there would likely be an important need to remove excess or poorly/non-functioning GPR63 from the cilium, a job which could very well require the retrograde transport capabilities of TTC21B. Future work could determine localization of GPR63 with either amino acid at position 407. In contrast, the entire RxR ER retrieval domain of GPR63 is lost in the *Gpr63^Del^* allele, suggesting a more profound effect in intracellular GPR63 trafficking in these animals.

### The role of Gpr63 in GLI3 processing

Our previous data has shown that GLI3 protein processing is disrupted in *Ttc21b^aln/aln^* mutants (Tran et al. 2008) and further unpublished data suggests this is indeed a substantial explanation for the embryonic phenotypes. We therefore hypothesize that reduced *Gpr63* function further impairs GLI3 activity in the *Ttc21b^aln/aln^* mutant. The GPCRs *Gpr161* and *Gpr175* have both been implicated in regulating GLI3 processing by controlling cAMP levels and PKA activity (Mukhopadhyay et al. 2013; Singh et al. 2015). *Gpr161* is present in the cilium prior to SHH stimulation, promoting GLI3 processing into GLI3R, while *Gpr175* enters after SHH stimulation to inhibit this process (Mukhopadhyay et al. 2013; Singh et al. 2015). One possibility is that *Gpr63* functions similarly and, like *Gpr175*, may be a context dependent modulator of SHH signaling, rather than an essential core component of the pathway. This is an intriguing idea that could potentially explain how loss of *Gpr63* may have tissue specific requirements for modulating SHH signaling.

## MATERIALS AND METHODS

### Animal husbandry

All animals were housed under an approved protocol in standard conditions. All euthanasia and subsequent embryo or organ harvests were preceded by Isoflurane sedation. Euthanasia was accomplished via dislocation of the cervical vertebrae. For embryo collections, noon of the day of vaginal plug detection was designated as E0.5. The *Ttc21b^aln^* allele used in this study has been previously published (Tran et al. 2008). *Ttc21b^aln^* mice were serially backcrossed to C57BL/6J (JAX:000664) and FVB/NJ (JAX:001800) mice to produce strain specific *Ttc21b^aln^* mice. Genotyping was performed by PCR, Sanger Sequencing, or Taqman assays (Table S4).

### QTL Analysis

GigaMUGA and MegaMUGA genotyping of recombinants was performed by Neogen (Morgan et al. 2015). The argyle and R/qtl analysis of GigaMUGA SNP data and brain size was performed at CCHMC (Broman and Sen 2009; Morgan 2015). Coding differences of candidate genes were compared using the Mouse Genomes Project website from Wellcome Sanger Trust Institute (https://www.sanger.ac.uk/sanger/Mouse_SnpViewer/rel-1505).

### CRISPR-Cas9

Donor oligonucleotide and guide RNA vector were designed using Benchling (Benchling, 2016). sgRNA sequence: TCCCTGGTCACACAAGTCGA. sgRNA were validated using Benchling and CRISPRscan with the following scores: Benchling-48.5 and CRISPRscan-49 (Benchling, 2016) (Moreno-Mateos, et al., 2015). Donor oligonucleotide sequence (lower case letters represent coding change): CCATAATCGCTCGATATGTTTCAAGAGTTCGGTATTCACAACACCGTCCGATGTTCCCCACACACGTAGACGGCACTAGGGCGAATgCGTCGcCTTGTaTGcCCgGGGAGCCGTGGCAAGAACT. Donor oligonucleotide was supplied by Integrated DNA Technologies. sgRNA was synthesized and together with donor oligonucleotide injected into FVB mouse zygotes by CCHMC Transgenic Animal and Genome Editing Core.

### Cell Culture and Transfection

A pCMV6-GPR63-myc expression plasmid (Origene, Rockville, MD) was transfected into mIMCD cells at ~80% confluency using Lipofectamine 3000 and incubated for 2-3 days. To induce ciliagenesis, cells were grown to confluency and serum was not changed over the course of incubation to induce starvation. Immunocytochemistry was performed with a Cells were stained with a rabbit anti-myc (Abcam ab9106) and anti-mouse acetylated-tubulin (SIGMA T6793) antibodies to detect cilia and Gpr63 localization. Secondary antibodies were Alexa Fluor-488 Goat anti-mouse/rabbit (Invitrogen, A11001, A20980). Confocal imaging was performed on a Nikon C2 system.

## ACKNOWLEDGEMENTS

Funding for this project comes from the National Institute of General Medical Sciences (R01 GM112744 R.W.S.).

